# Extracellular loops of the β-barrel domain catalyze rapid folding for function of self-associating autotransporters

**DOI:** 10.1101/2025.04.08.647479

**Authors:** Xiaojun Yuan, Matthew D. Johnson, Jiansi Long, Jing Zhang, Alvin W. Lo, Jason J. Paxman, Duy Phan, Begoña Heras, Mark A. Schembri, Gerard H. M. Huysmans, Ian R. Henderson, Matthew T. Doyle, Denisse L. Leyton

## Abstract

Bacterial aggregation is a phenotype associated with disease pathogenesis. Aggregate formation enhances biofilm development, host colonization, and resistance to antibiotics and host defenses. Antigen 43 (Ag43) is a surface-located autotransporter produced by pathogenic *Escherichia coli* that mediates cell aggregation in biofilms. Two Ag43 molecules, each from neighboring bacterial cells, fold into elongated β-helical passenger domains that associate in a head-to-tail manner while being anchored to the cell surface by their outer membrane-embedded β-barrels. In this study, we conduct mutational analyses on Ag43 to show that the β-hairpin structure of the fourth and fifth extracellular loops of the β-barrel domain have a crucial role for passenger domain folding and subsequent formation of bacterial aggregates. This work provides mechanistic insight into the role of the autotransporter β-barrel domain to nucleate the rapid folding of the passenger domain into the β-helix that enables bacterial interactions during infection.

## Introduction

“Classical” autotransporters are a superfamily of bacterial virulence factors that are produced by most Gram-negative bacteria^1^. The classical autotransporters can be grouped into 16 distinct subfamilies based on amino acid identity where those within each subfamily have similar structural and functionals features^2^. Adhesion, invasion, aggregation, biofilm formation, toxicity, and immune evasion are attributes of many different autotransporters^3^. Some proteins can mediate more than one of these functions. The autotransporters share a common modular organization of a 12-stranded β-barrel domain embedded in the outer membrane, forming part of a secretion pore that is needed to translocate the covalently linked passenger domain to the bacterial cell surface^4–8^. Passenger translocation occurs in a C- to- N-terminal direction, where a conserved structural element called the autochaperone (AC) domain appears first at the cell surface^9–13^. Structures of AC domains show that their C-termini comprise of three β-strand rungs capped by a β-hairpin formed by the last two anti-parallel β-strands of the AC domain^14^. Several studies have shown that this domain is required for folding of the elongated β-helical structure^15–17^ that is common to most autotransporters and that mediates the biological function(s) of the protein^1,9,18,19^.

In a cellular context, key molecules catalyze the folding and membrane insertion stages for classical autotransporter β-barrels, as they do for other types of outer membrane β-barrel proteins (OMPs)^7,8,10,20–27^. For example, recent evidence from *in vivo* cross-linking and single-particle cryogenic electron microscopy studies showed that BamA, the essential component of the β-barrel assembly machinery^28–30^, mediates the integration of a partially folded EspP β-barrel from the periplasm into the outer membrane via the formation of a hybrid barrel^8,27,31^. Furthermore, a separate study used *in vivo* cross-linking to show that the EspP passenger domain is translocated through the pore formed by the transient fusion of the BamA-EspP barrels during membrane insertion^7^. However, we and others have shown that intrinsic features of the autotransporter β-barrel domain also enforce control over the assembly process. Point mutations resulting in the modest distortion of the Pet and EspP β-barrel scaffolds were shown to slow the rate of passenger translocation^26,32^. Similar observations were made upon truncation of the fourth extracellular loop (L4) of the BrkA and Pet β-barrels^33,34^. More recently, folding of the Pet and EspP passenger domains into a β-helix were shown not to be spontaneous, but rather nucleated by the β-hairpin structure of the fifth extracellular loop (L5) of the β-barrel domain^34^. Given that the AC domain appears first at the bacterial cell surface, it was hypothesized that L5 transiently interacts with this region to initiate its folding into a stack of β-helical rungs on which the remainder of the N-terminal β-helical backbone is folded^34^.

This work provided significant new insights into the contribution of the β-barrel domain to the biogenesis of Pet and EspP, both of which are members of the SPATE (serine protease autotransporters of *Enterobacteriaceae*) subfamily. However, it is not known if L5-assisted folding of the passenger domain is a fundamental process widely conserved across autotransporters. Furthermore, the region of the passenger that interacts with L5 has not yet been identified, nor whether the loop constitutes a mechanism to accelerate folding and, therefore, enable autotransporter function in the absence of external energy inputs. To better understand the nature of passenger domain folding towards therapeutic strategies to block autotransporter function, we examined the role of the L5 and L4 regions on the autotransporter antigen 43a (Ag43a). Ag43a is a prototype member of the self-associating autotransporters (SAAT) subfamily (divergent from SPATEs) that is produced by most *Escherichia coli* pathotypes and that promotes bacterial aggregation and biofilm formation by Ag43a-Ag43a binding between adjacent cells^14,35–38^.

The experiments described here provide evidence that the Ag43a β-barrel domain actively accelerates the rate of folding of the passenger domain. Our work suggests that it is the β-hairpin structure of the long extracellular loops of the β-barrel domain that provide a structural template for the rapid nucleation of passenger folding into a stable β-helix. Biophysical studies on truncated L4 and/or L5 β-hairpins show that the accelerated folding of the passenger domain can occur independently of BamA and the other cellular machinery when refolded *in vitro*. Furthermore, by monitoring the assembly of Ag43a mutants with truncated L4 and/or L5 β-hairpins in live bacterial cells, we demonstrate that rapid loop-mediated folding of the passenger domain is crucial for the biological function of this SAAT.

## Results

### Cleavage between α^43^ and β^43^ is autocatalytic

Once assembled correctly into the outer membrane, Ag43a is cleaved by a suspected autocatalytic reaction that occurs within the passenger between D^552^ and P^553^, resulting in the release of the functional domain (α^43^) from the translocation domain (β^43^), where β^43^ encompasses the AC and β-barrel domains^36,39^. Thus, the AC domain remains attached to the β-barrel domain despite the former being an extension of the passenger β-helix (that is capped by a β-hairpin) as shown in the crystal structure of an unprocessed mutant of Ag43 from UPEC UTI89 where α^43^ and the AC domain are covalently linked^14^. There is no available structure for the Ag43a β-barrel domain. Therefore, to map the locations of L4 and L5 in the Ag43a β-barrel domain, we used AlphaFold3^40^ to generate structural (Fig. 1a) and topology (Fig. 1b) models of the β^43^ domain. The models suggest that L4 (residues D^900^ – H^913^) and L5 (S^969^ – N^992^) interact with each other forming a four-stranded β-sheet. To investigate if L4, L5, and/or L4 + L5 are required for translocation and/or folding of the Ag43a passenger domain, we constructed Ag43^Δ1-206^ and derivates containing deletions of residues D^900^ to F^911^ and/or R^971^ to S^990^ (Figs. 1c and d). Given that the N-terminal truncation of the passenger domain (residues 1-206) removes most of the stem of the L-shape responsible for the Ag43a self-association mechanism^36^, we reasoned that this truncation would prevent the formation of aggregates to enable accurate monitoring of protein folding *in vitro*.

**Figure 1.**
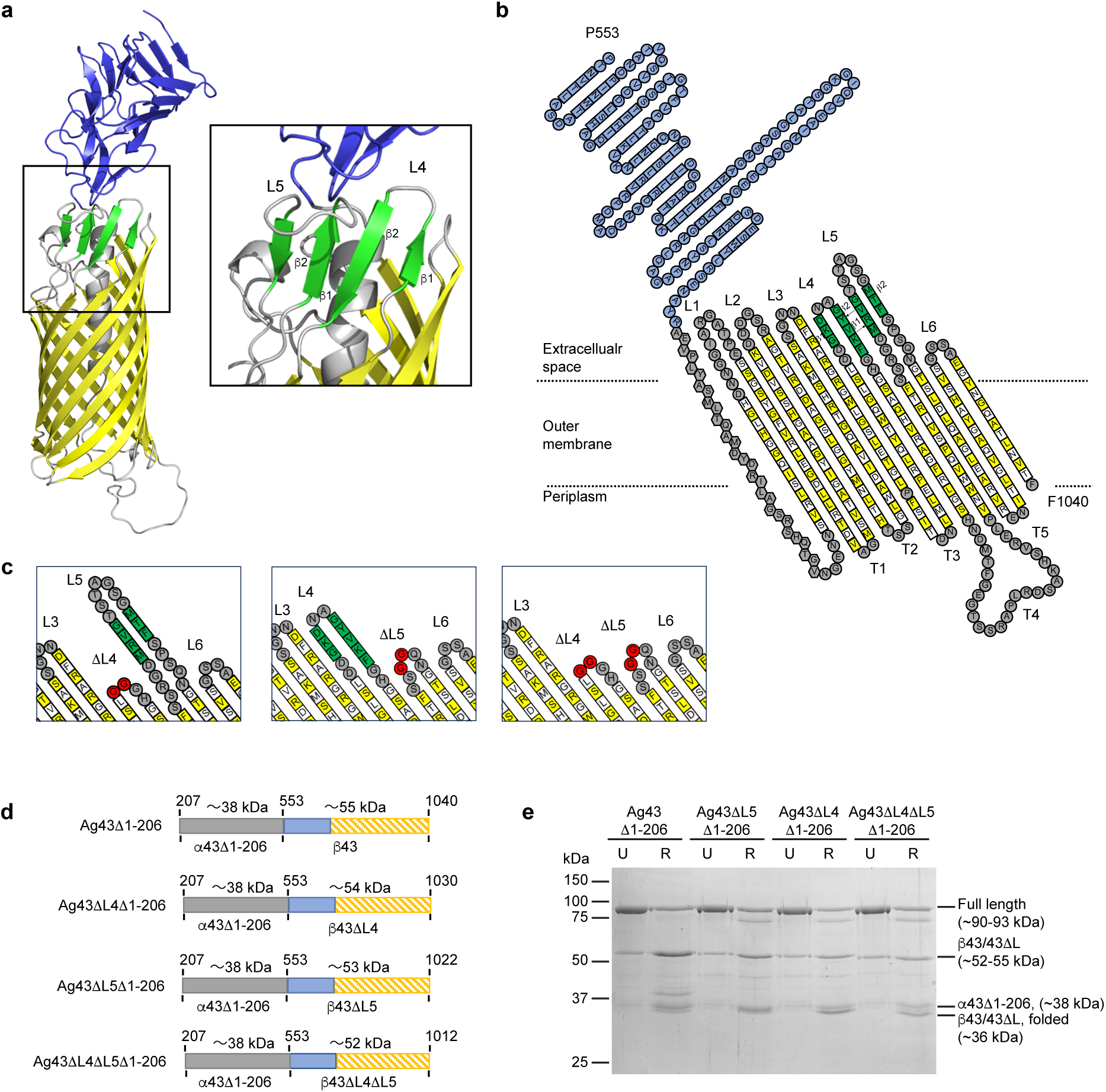
Autocatalysis between α^43^ and β^43^ *in vitro* mimics the *in vivo* cleavage reaction. AlphaFold3 (**a**) and topology (**b**) models of the β^43^ domain. The topology model of β^43^ is based on **a**. For **a**, β-strands 1 – 12 of the β-barrel are shown in yellow and for **b**, white and yellow squares indicate residues with side chains that point toward the β-barrel lumen and lipid bilayer, respectively. Residues in the α-helical segment, periplasmic turns (labeled T1-T5 in topology model), and extracellular loops (labeled L1, L2, L3, and L6 in topology model) are shown in grey. Residues in extracellular loops labeled L4 and L5, and in the AC are shown in green and blue, respectively. **c** Topology models of β^43^ showing the L4 truncation created by replacing D^900^ to F^911^ with 2 glycine residues (shown in red) (left panel), the L5 truncation created by replacing R^971^ to S^990^ with 2 glycine residues (shown in red) (middle panel), and the double L4 + L5 truncations created as per the single deletions (right panel). **d** Schematic of the Ag43^Δ1-206^, Ag43ΔL4^Δ1-206^, Ag43ΔL5^Δ1-206^, and Ag43ΔL4ΔL5^Δ1-206^ proteins. The 346-residue passenger incorporates residues 207 to 552 of Ag43a (shown in grey) immediately upstream of the wild-type (488 residues; 553 to 1040), L4-truncated (478 residues; 553 to 1030), L5-truncated (470 residues; 553 to 1022), or L4 + L5-truncated (460 residues; 553 to 1012) β^43^ domains. **e** Refolding of Ag43^Δ1-206^, Ag43ΔL4^Δ1-206^, Ag43ΔL5^Δ1-206^, and Ag43ΔL4ΔL5^Δ1-206^ was monitored by SDS-PAGE and Coomassie Brilliant Blue staining of the gels. Processing of the passenger domain resulted in the autocatalytic conversion of the ∼93 kDa full-length protein to ∼38 kDa and ∼52-55 kDa proteins. U, unfolded protein; R, refolded protein. Refolding of these proteins also visualized an ∼80 kDa species that is absent in Ag43^Δ1-206^ and an ∼40 kDa species that is present only in Ag43^Δ1-206^. These species correspond to a N-terminally truncated passenger domain still attached to its translocation domain and an additional folded β^43^ species that has been observed previously^41^, respectively.

To test the hypothesis that cleavage at the α^43^–β^43^ junction between residues D^552^ and P^553^ is autocatalytic, urea solubilized Ag43^Δ1-206^, Ag43ΔL4^Δ1-206^, Ag43ΔL5^Δ1-206^, or Ag43ΔL4ΔL5^Δ1-206^ were refolded by rapid dilution into LDAO detergent, as described previously^32,34^, and the accumulation of ∼52-55 kDa and ∼38 kDa cleavage products corresponding to β^43^ and α^43Δ1-206^ was monitored by SDS-PAGE (Fig. 1e). The minor size differences between β^43^, β^43^ΔL4, β^43^ΔL5, and β^43^ΔL4ΔL5 reflect the decrease in size of the mutant proteins after loop truncation. This contrasts with the α^43Δ1-206^ domains, which are all the same molecular weight because these ∼38 kDa species are no longer attached to their respective β^43^ domains. Most folded β-barrel proteins are resistant to denaturation by SDS at low temperatures, migrating faster during electrophoresis than when unfolded^42^. As shown previously^41,43^, β^43^ is unusual as it is partly resistant to denaturation even at 100 °C; the boiling of samples was not sufficient to completely denature any of the cleaved translocation domains (detected using anti-Ag43a β-barrel antibody), resulting in the species migrating near the 37 kDa standard (Fig. 1e).

N-terminal sequencing revealed that the ∼55 kDa β^43^ fragment starts with the cleavage site sequence (P^553^TNVT) (Supplementary Fig. 1a). These data show that the refolded proteins are processed between D^552^ and P^553^ in a manner that mimics the *in vivo* cleavage reaction. Similar observations were made previously for the *E. coli* SAAT adhesin involved in diffuse adherence (AIDA-I), an Ag43a homologue. Processing of the AIDA-I (passenger) domain from the AIDAc (translocation) domain is also autocatalytic^44^. Even with the proteolytic processing, in bacterial cells, α^43^ and β^43^ remain associated through non-covalent interactions that can be disrupted by brief heating to 60 °C to release α^43^ from the cell surface^45^. To determine if the ∼52-55 kDa β^43^ and ∼38 kDa α^43Δ1-206^ cleavage products resulting from autocatalytic processing of the full-length ∼90-93 kDa proteins remained associated post refolding, we purified the proteins from the refolding buffer via his-tag affinity. This showed that the α^43Δ1-206^ fragments co-eluted with the β^43^ fragment (Supplementary Fig. 2), indicating that α^43^ and β^43^ remain associated after refolding and autocleavage as seen *in vivo*^45^.

### L4 and L5 accelerate folding of the Ag43a passenger and AC domains *in vitro*

To further measure the role of L4 and L5 on passenger folding, urea denatured Ag43^Δ1-206^ and loop-truncated derivatives were subjected to refolding by rapid dilution out of the denaturant. During refolding, timepoint samples were taken and subjected to trypsin treatment to monitor the appearance of fragments corresponding to protease-resistant folded segments^34^. The samples, probed using an anti-α^43^ antibody, showed that the passenger (∼38 kDa) released from wild-type β^43^ was largely resistant to trypsinolysis, generating one dominant fragment of ∼33 kDa by as early as 5 min (Fig. 2a, top panel). There was a delay in the release of folded passenger from the ΔL4 and L5 mutants (Fig. 2b,c, top panel, 45 and 10 min, respectively), and the ΔL4ΔL5 mutant released barely detectable levels of folded passenger by 4 h (Fig. 2d, top panel). Given that the yield of total released passenger from β^43^, β^43^ΔL4, β^43^ΔL5, and β^43^ΔL4ΔL5 by 4 h is similar in the absence of trypsin (Fig. 2, top panel, −trypsin), these data suggest that L4 and L5 within the Ag43a translocation domain are required for efficient folding of the passenger domain *in vitro*. These data also show that adoption of the fully folded state of the passenger domain is not necessary for its autocatalytic release from the translocation domain. The samples were also probed with anti-Ag43a β-barrel antibody. While trypsin-resistant fragments between ∼15-30 kDa were observed at much higher levels for the loop-truncated mutants than the wild-type translocation domains, they were generated by 0 min in all cases (Fig. 2, bottom panel). Our previous work on Pet showed that truncation of L5 does not inhibit or delay folding of the β-barrel domain *in vitro*, resulting in the accumulation of trypsin-resistant fragments corresponding to the L5-truncated β-barrel by 0 min^34^. Therefore, we propose that these low-molecular weight species result from trypsin digestion of the wild-type and loop-truncated Ag43a β-barrel domains. Consequently, we propose that the relatively late accumulation of trypsin-resistant fragments between ∼34-44 kDa for the loop-truncated mutants results from digestion of the slow to fold AC domains (Fig. 2, bottom panel).

**Figure 2.**
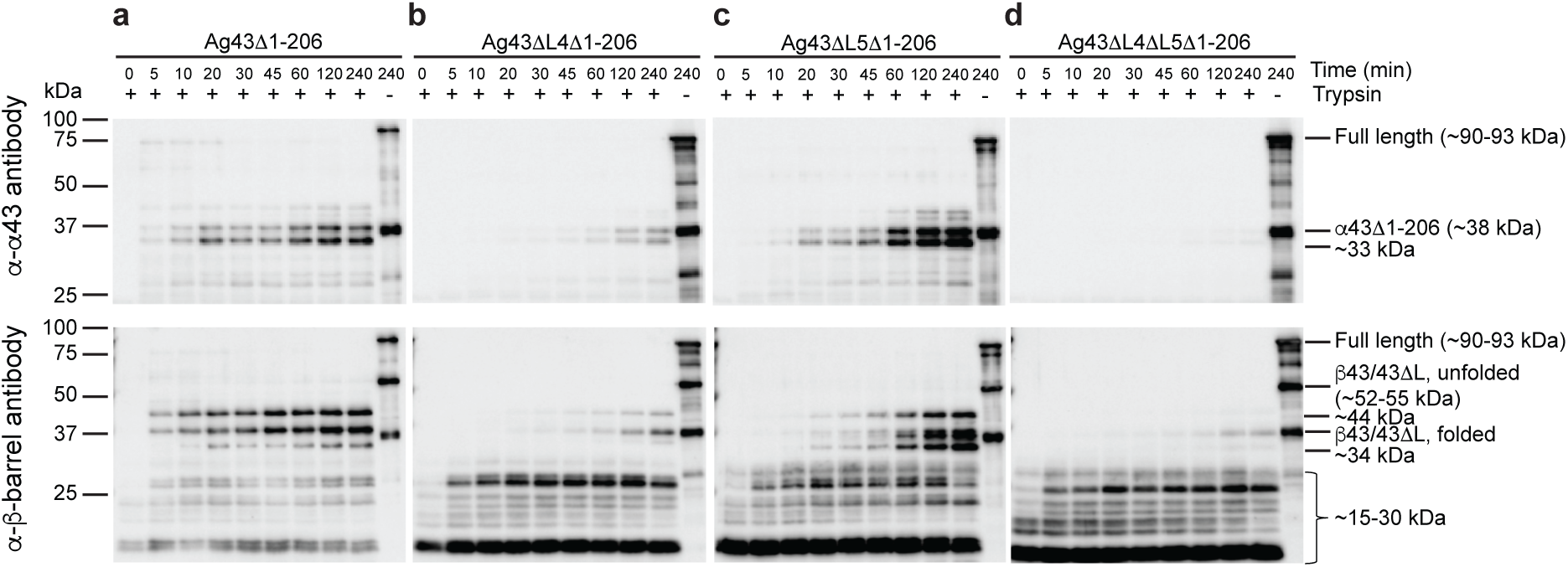
L4 and L5 accelerate folding of the Ag43a AC and passenger domains *in vitro*. Folding of Ag43^Δ1-206^ (**a**), Ag43ΔL4^Δ1-206^ (**b**), Ag43ΔL5^Δ1-206^ (**c**), and Ag43ΔL4ΔL5^Δ1-206^ (**d**) monitored by the accumulation of trypsin-resistant fragments. Samples were probed by Western immunoblotting using anti-α^43^ (top panels) and anti-β-barrel (bottom panels) antibodies.

To validate that the loop truncations do not affect β-barrel-assembly, we also purified urea solubilized Ag43^Δ1-705^ and loop-truncated derivatives comprising of only the β-barrel domains and refolding was examined by a time course trypsin assay as described above. All forms of the protein resolved into a similar set of tryptic products by 0 min (Fig. 3a), suggesting that truncation of L4 and/or L5 does not slow folding of the Ag43a β-barrel domain *in vitro*. To test this hypothesis further, we monitored folding of the wild-type and ΔL4ΔL5 β-barrel domains by far-UV circular dichroism (CD) and tryptophan fluorescence spectroscopy. The CD spectra of both refolded β-barrels showed a single minimum of ellipticity at ∼217 nm, a characteristic of β-sheets (Fig. 3b). No significant differences were observed between the spectra taken at 0 and 4 h, showing that both β-barrels acquired secondary structure immediately after rapid dilution of the denaturant. Tryptophan fluorescence spectra also showed rapid folding of both β-barrels with blue-shifted spectra immediately observed upon urea dilution (Fig. 3c). Moreover, a melt-curve was generated by measuring ellipticity at 218 nm at increasing temperatures, which showed that the thermal stability of both β-barrels is similar (Fig. 3d). These data strongly suggest that the delayed folding of the Ag43a passenger and AC domains by the ΔL4, ΔL5, or ΔL4ΔL5 β-barrels is directly related to the function of the loops rather than defective β-barrel folding or stability.

**Figure 3.**
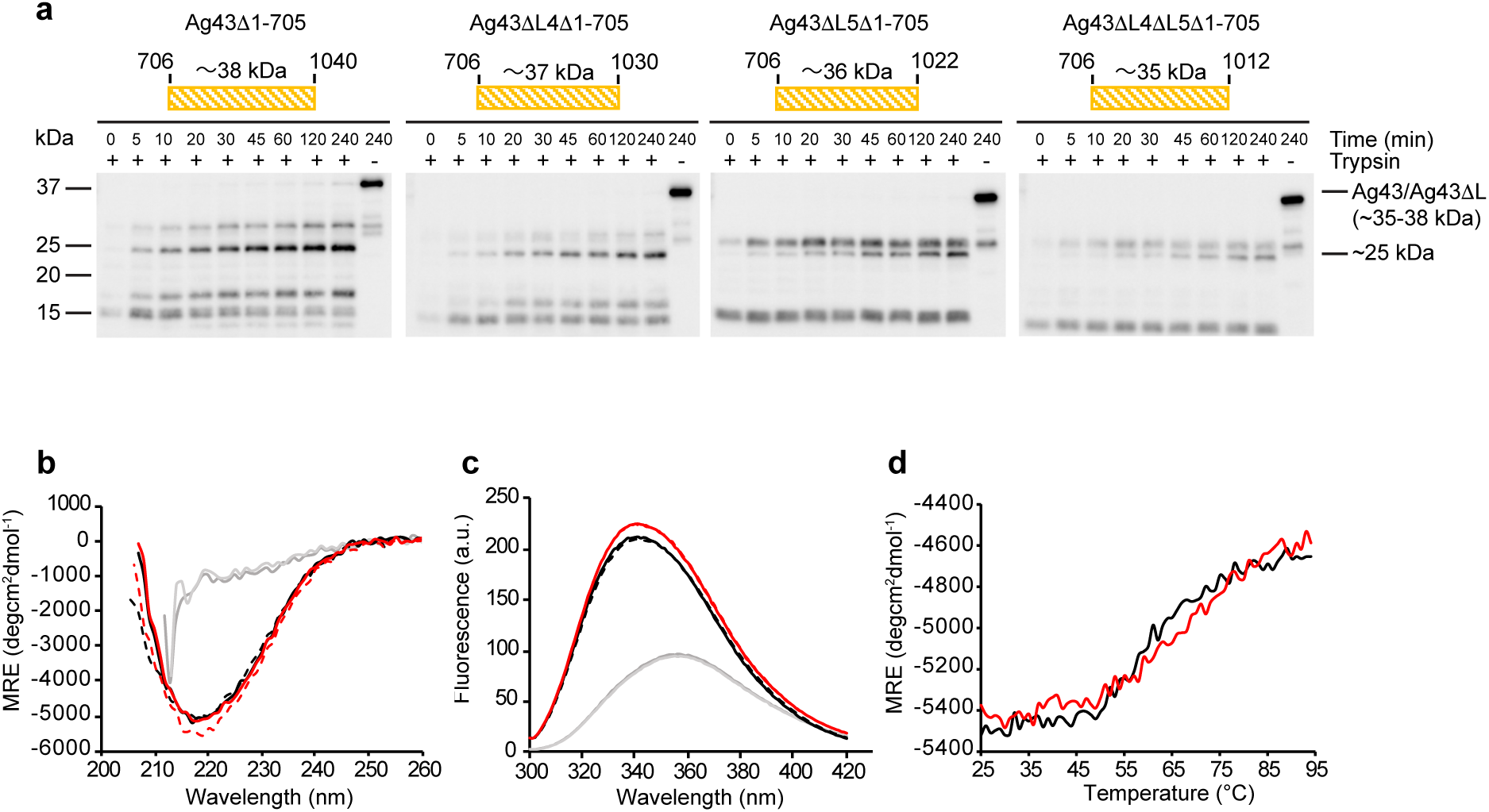
Truncations of L4 and L5 do not slow folding of the Ag43a β-barrel *in vitro*. **a** Schematic of the Ag43^Δ1-705^ (top, first panel from left), Ag43ΔL4^Δ1-705^ (top, second panel from left), Ag43ΔL5^Δ1-705^ (top, third panel from left), and Ag43ΔL4ΔL5^Δ1-705^ (top, fourth panel from left) proteins comprising the wild-type and loop-truncated β-barrels only. Folding of these β-barrels was monitored by the accumulation of trypsin-resistant fragments (bottom panels) as in Figure 2 and probing with anti-β-barrel antibody. Far-UV CD **(b)** and tryptophan fluorescence **(c)** spectra of unfolded protein (grey) and *in vitro* folded Ag43^Δ1-705^ (black) and Ag43ΔL4ΔL5^Δ1-705^ (red) β-barrel domains at 0 min (dashed lines) or 4 h (solid lines). **d** Thermal denaturation of *in vitro* folded Ag43^Δ1-705^ (black) and Ag43ΔL4ΔL5^Δ1-705^ (red) β-barrel domains monitored at 218 nm. MRE; mean residue ellipticity.

### Truncations of L4 and/or L5 perturb folding of the Ag43a passenger and AC domains *in vivo*

To assess whether the delayed folding observed for the loop truncation mutants *in vitro* impacted on autotransporter assembly in bacterial cells, we produced Ag43a or the loop-truncated mutants in the previously described *E. coli ag43* null strain MS427^46^ and monitored surface folding of the passenger by subjecting the bacteria to proteolysis by proteinase K. Western immunoblotting of lysates for the passenger demonstrated the presence of the full-length Ag43 (∼109 kDa), Ag43ΔL4 (∼108 kDa), Ag43ΔL5 (∼107 kDa), and Ag43ΔL4ΔL5 (∼106 kDa) pro-proteins, and an ∼55 kDa species corresponding to the α^43^ passenger segment (Fig. 4a, top left panel, +OmpT, −/+PK). Curiously, comparison of the assembly of the loop-truncated pro-proteins revealed the presence of a ∼66 kDa passenger fragment, which was partially digested by proteinase K into ∼55 kDa and ∼40 kDa protease-resistant fragments (Fig. 4a, top left panel, +OmpT, compare −/+PK). Proteinase K proteolysis of Ag43ΔL4ΔL5-AB, an Ag43ΔL4ΔL5 derivative containing a mutation in the autocatalytic cleavage site (D^552^A) that results in translocation of the passenger domain to the cell surface but not its autocatalysis or heat-released from the cell, confirmed that the ∼66 kDa passenger fragment gives rise to both protease-resistant fragments (Supplementary Fig. 3, +PK, compare Ag43ΔL4ΔL5 with Ag43ΔL4ΔL5-AB). The adduct of the ∼55 kDa and ∼40 kDa protease-resistant fragments exceeding 66 kDa suggests that digestion of the ∼66 kDa passenger fragment first generates a ∼55 kDa protease-resistant fragment that is subsequently digested into a ∼40 kDa protease-resistant fragment. Furthermore, the lower relative abundance of these protease-resistant fragments compared to the ∼66 kDa passenger fragment suggests that a population of the ∼66 kDa molecules are unfolded and completely degraded by the enzyme.

**Figure 4.**
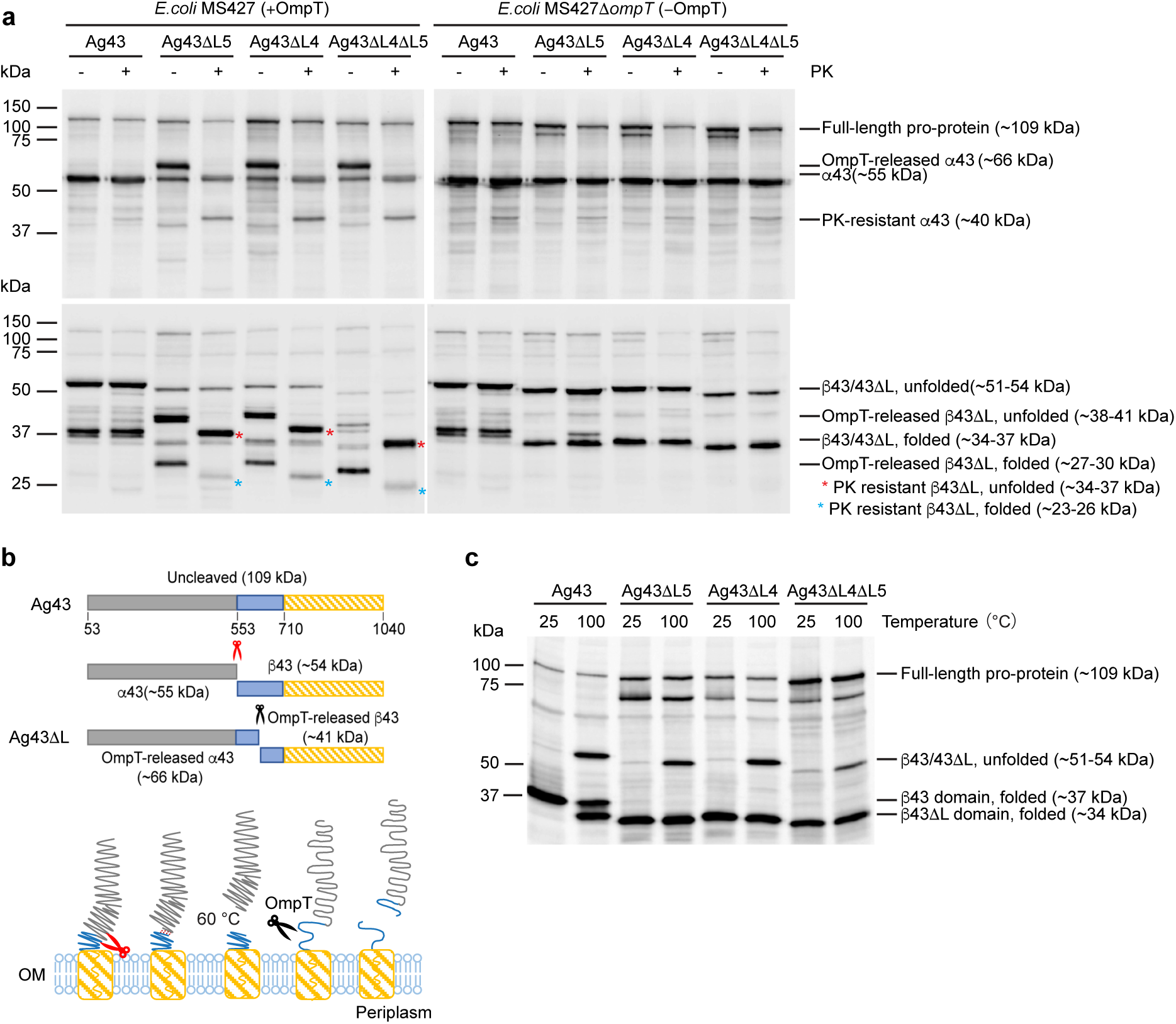
Truncations of L4 and L5 perturb folding of the Ag43a AC and passenger domains in bacterial cells. **a** Expression of Ag43, Ag43ΔL4, Ag43ΔL5, and Ag43ΔL4ΔL5, and sensitivity to proteinase K (PK) in *E. coli* MS427 (left panels) and *E. coli* MS427Δ*ompT* (right panels) monitored by Western immunoblotting with anti-α^43^ antibody (top panels) or anti-β-barrel domain antibody (bottom panels). Expression of the mutant proteins in MS427Δ*ompT* also visualized an ∼80 kDa species that is absent in wild-type Ag43 in **a** and **c** (as in Figure 1e) corresponding to a species susceptible to OmpT degradation. **b** The domain structure (top) and model for Ag43 biogenesis (bottom). Post insertion of β^43^ into the outer membrane (OM) and translocation of α^43^ onto the cell surface, a cleavage reaction at the α^43^-β^43^ junction (red scissors) releases the ∼55 kDa α^43^ domain that remains non-covalently bound to the surface via β^43^ (∼54 kDa). The α^43^ domain can be released by brief heat treatment. In the absence of L4 and/or L5, the unfolded AC domain is susceptible to cleavage by OmpT, which liberates a ∼66 kDa α^43^ fragment into the extracellular milieu. **c** Total membranes from Ag43, Ag43ΔL4, Ag43ΔL5, and Ag43ΔL4ΔL5 expressing cells were heated at the temperatures indicated and analyzed by Western immunoblotting with anti-β-barrel antibody.

### Truncations of L4 and/or L5 expose a cryptic OmpT cleavage site in the Ag43a AC domain

Our *in vivo* data suggest that proteinase K digestion of the ∼66 kDa passenger fragment gives rise to a population of folded ∼55 kDa protease-resistant molecules through digestion of a ∼11 kDa region that is completely degraded by the enzyme and is, therefore, presumably unfolded. Since OmpT (an outer membrane protease endogenous to *E. coli*) has been shown to digest unfolded passenger domains on the bacterial cell surface, while surface exposed and natively folded passenger domains are resistant to OmpT degradation^16,34^, limited proteinase K proteolysis of Ag43, Ag43ΔL4, Ag43ΔL5, and Ag43ΔL4ΔL5 was repeated in *E. coli* MS427Δ*ompT*, a strain lacking OmpT. The limited proteinase K proteolysis assay revealed that the ∼66 kDa passenger fragment is absent even in the absence of proteinase K (Fig. 4a, top right panel, −OmpT, −PK), thereby showing that this fragment is the product of OmpT digestion. Furthermore, the mostly comparable steady state, relative abundance of the ∼55 kDa α^43^ functional domains and ∼40 kDa protease-resistant fragments in all samples suggests that in the absence of OmpT, most of the passenger and AC domain molecules secreted by β^43^ΔL4, β^43^ΔL5, and β^43^ΔL4ΔL5 fold into proteinase K-resistant conformations by the time they are subjected to proteolysis (*t*= 2 h) (Fig. 4a, top right panel, −OmpT, +PK). This supports of our *in vitro* data that suggests that in the absence of L4 or L5, the folding of the passenger and AC domain by the β-barrel is delayed but not completely inhibited (Fig. 2). To understand the identity of the OmpT-released ∼66 kDa passenger fragment, we isolated it from culture supernatants where it accumulated when cells expressed the loop-truncated mutants (Supplementary Fig. 4). N-terminal sequencing revealed that the fragment from the Ag43ΔL5 culture supernatant sample begins with (A^53^DIVV) (Supplementary Fig. 1b), which corresponds to the residues of the passenger domain immediately after signal sequence removal. Since this fragment is larger than the ∼55 kDa α^43^ functional domain, together these data suggest that when any of the loops are truncated, OmpT has access to an unfolded region downstream of the passenger cleavage site, located in the AC domain (Fig. 4b). Furthermore, since the OmpT-released passenger fragment is ∼66 kDa and not ∼11 kDa, these data suggest that OmpT digests the unfolded AC domain prior to autocatalysis of α^43^ from β^43^.

The samples were also probed with anti-Ag43a β-barrel antibody to test the effect of the loop truncations on folding of the Ag43a β-barrel in bacteria. The presence of cleaved β^43^ species in unfolded (∼51−54 kDa) and folded (∼34−37 kDa) forms (Fig. 4a, bottom left panel, +OmpT, −/+PK) suggested that autocatalysis of the loop-truncated pro-proteins, like in wild-type Ag43, occurred at the α^43^−β^43^ junction (Fig. 4b). The cleaved β^43^ΔL4, β^43^ΔL5, and β^43^ΔL4ΔL5 species were present in much reduced quantities relative to wild-type β^43^ (Fig. 4a, bottom left panel, +OmpT, −/+PK) and to the equivalent samples assayed in MS427Δ*ompT* (Fig. 4a, bottom panels, compare +OmpT [left panel] versus −OmpT [right panel], −/+PK). This is likely due to there being less of the loop-truncated pro-proteins relative to the wild-type pro-protein available for autocatalysis due to OmpT-mediated release of the ∼66 kDa passenger fragment. The loop-truncated pro-proteins generated additional cleaved β^43^ species in unfolded (∼38−41 kDa) and folded (∼27−30 kDa) forms only in MS427 (Fig. 4a, bottom panels, compare +OmpT [left panel] versus −OmpT [right panel], −PK) that were converted into smaller species after proteinase K treatment (Fig. 4a, bottom left panel, +OmpT, compare −PK versus +PK). This suggests that the ∼38−41 kDa fragments correspond to what remain of the ΔL4, ΔL5, and ΔL4ΔL5 translocation domains following OmpT release of the ∼66 kDa passenger fragment. To compare the thermostability of β^43^ and the loop-truncated translocation domains, and their ability to fold into the outer membrane *in vivo*, we probed unheated and heated samples by anti-β-barrel Western blot. All the β-barrels showed greater electrophoretic mobility in samples heated at 25 °C and with partial denaturation of the β-structure at 100 °C (Fig. 4c). Together, these data show that none of the loop truncations inhibit correct assembly of the β-barrel into the outer membrane.

### Passenger and AC domain folding relies on the β-hairpin conformation of L5

Our previous work showed that a stretch of six amino acids in the first β-strand of L5 forms a template for the folding of the Pet passenger domain in a manner that is reminiscent of β-strand augmentation^34^. To determine if the first and/or second β-strand in L5 are responsible for mediating folding of the Ag43a passenger and AC domains, Ag43a variants were created (Fig. 5a) and analyzed by monitoring the appearance of the ∼66 kDa OmpT-released α^43^ fragment in *E. coli* MS427 and MS427Δ*ompT* since an AC domain that is unable to efficiently fold into its native conformation will be susceptible to cleavage by OmpT. Ag43L5β1G, a variant with most amino acids in the first β-strand in L5 (D^973^-V^976^) mutated to glycine demonstrated a severe defect in AC domain folding like that observed for Ag43ΔL5, evidenced by the appearance of the ∼66 kDa OmpT-released α^43^ fragment in *E. coli* MS427, but not in MS427Δ*ompT* (Fig. 5b). Ag43L5β2/G, a variant with most amino acids in the second β-strand in L5 (T^986^-P^989^) mutated to glycine showed a similar phenotype, except that less of the ∼66 kDa OmpT-released α^43^ fragment was found in the culture supernatant, and there was more of the ∼55 kDa α^43^ surface-bound fragment resulting from cleavage of the Ag43L5β2G pro-protein (Fig. 5b). Since a flexible cluster of glycine residues is a well-known secondary structure breaker^47^, we propose that substitution of the residues in the first and second β-strands in L5 with a stretch of glycine residues severely perturbs folding of the AC domain through disruption of the β-sheet structure in these regions.

**Figure 5.**
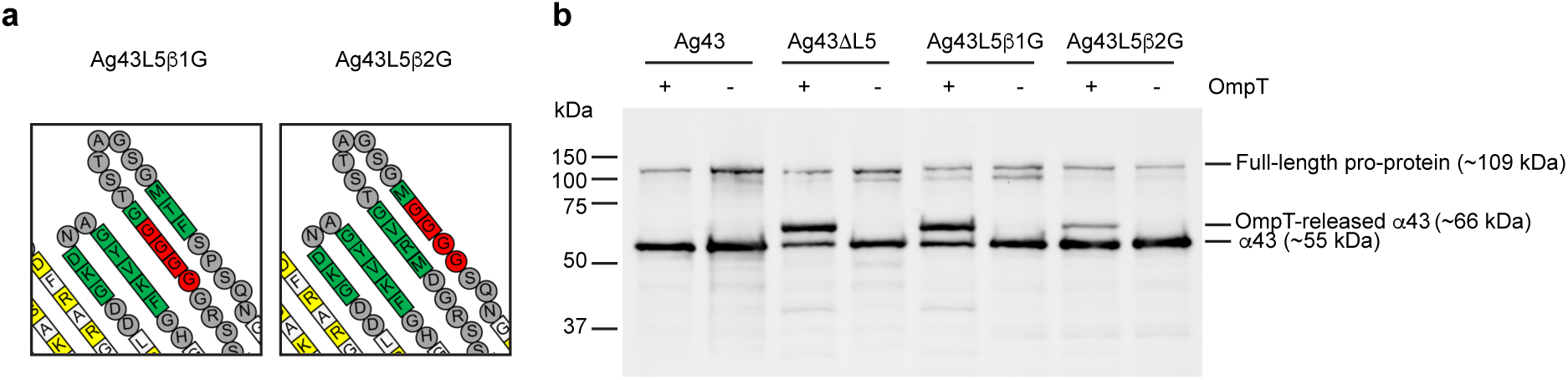
L5 likely templates folding of the Ag43 AC domain by β-strand augmentation. **a** Topology model of β^43^ showing the Ag43L5β1G (left panel) and Ag43L5β2G (right panel) mutations (in red). **b** Expression of Ag43, Ag43ΔL5, Ag43L5β1G, and Ag43L5β2G, and sensitivity to OmpT in *E. coli* MS427 (+OmpT) and *E. coli* MS427Δ*ompT* (−OmpT) monitored by Western immunoblotting with anti-α^43^ antibody.

### Ag43a-mediated bacterial clumping relies on L4 and/or L5

*E. coli* MS427 expressing Ag43, Ag43ΔL4, Ag43ΔL5, or Ag43ΔL4ΔL5 were examined in cell aggregation assays to test if OmpT-mediated release of the ∼66 kDa α^43^ fragments altered the proteins’ function. Our findings indicate that truncation of L4 and/or L5 abolished Ag43-mediated cell aggregation in all three mutant strains, which had an aggregation profile almost indistinguishable from that of the empty vector control, a strain completely lacking Ag43 (Fig. 6a, *E. coli* MS427 [+OmpT]). In contrast, when we repeated the cell aggregation assay in MS427Δ*ompT*, the ability of bacteria expressing Ag43ΔL4 or Ag43ΔL5 to mediate cell aggregation was delayed, but eventually restored almost to wild type levels (Fig. 6b, *E. coli* MS427Δ*ompT* [−OmpT]). The opposite was observed for Ag43ΔL4ΔL5, which failed to significant aggregate in the presence or absence of OmpT (Fig. 6a,b, *E. coli* MS427 [+OmpT] versus MS427Δ*ompT* [−OmpT]). This suggests that in MS427Δ*ompT*, the AC domains secreted by the loop-truncated β-barrels are not cleaved by OmpT. Instead, the full-length pro-proteins are autocatalyzed into α^43^ and β^43^ fragments that remain associated through non-covalent interactions, where the α^43^ domains from Ag43ΔL4, Ag43ΔL5, and to a very minimal extent, Ag43ΔL4ΔL5 are eventually able to fold into their native conformations and mediate cell aggregation, despite the absence of the loops. These observations suggest that the inability of Ag43ΔL4ΔL5 to mediate significant aggregation of MS427Δ*ompT* and by extension, the delay in aggregation observed for MS427Δ*ompT* strains expressing Ag43ΔL4 or Ag43ΔL5 relative to that harboring wild type Ag43, is a consequence of delayed folding in a population of passenger molecules during the time-course of the aggregation assay.

**Figure 6.**
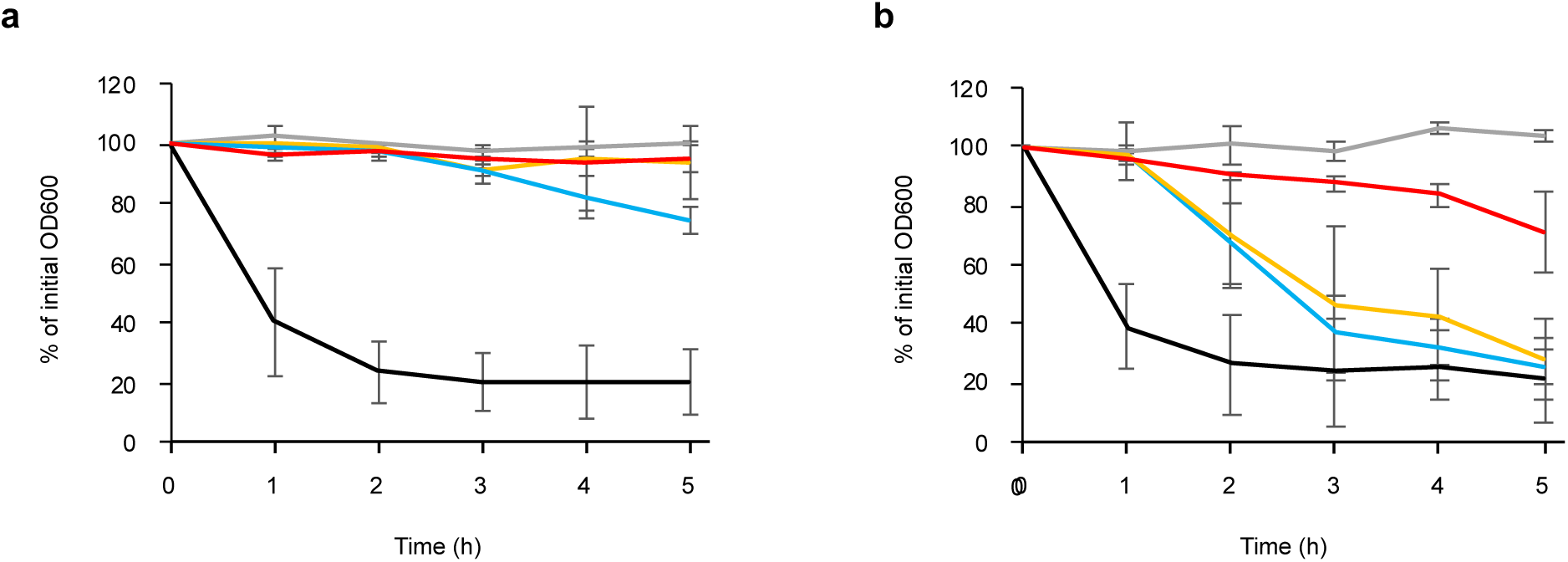
Ag43a-mediated bacterial clumping relies on L4 and/or L5. *E. coli* MS427 (+OmpT) **(a)** and *E. coli* MS427Δ*ompT* (−OmpT) **(b)** aggregation profiles of strains expressing Ag43 (black), Ag43ΔL4 (yellow), Ag43ΔL5 (blue), Ag43ΔL4ΔL5 (red), or empty vector (grey). In all cases, strains expressing the loop truncation mutants were compared with strains expressing Ag43 or empty vector. Shown are the means for at least three independent experiments.

## Discussion

In this current study, we analyzed the folding of Ag43 *in vitro* and in bacterial cells to test if loop-assisted folding of the passenger domain is conserved beyond the SPATE subfamily of autotransporters. Importantly, we also examined the influence of loop truncations on Ag43-mediated cell aggregation to test if the extracellular loops influence autotransporter biological function. We find that both L4 and L5 are required to rapidly fold the Ag43 AC and α^43^ domains into their native, protease-resistant structures in a biologically relevant time-scale, and that these long extracellular loops are indirectly required to mediate the virulence function of this SAAT protein. In a cellular context, this process is compatible with BamA-mediated integration of the autotransporter β-barrel domain into bacterial outer membranes^8^ and with subsequent translocation of the autotransporter passenger domain through a hybrid barrel^7^. We speculate that BamA is unlikely to be involved in autotransporter β-barrel-mediated folding of the passenger domain as we find that this process occurs independently of BamA (and other cellular machinery) *in vitro*.

We show that the amino acids of the passenger that interact with L4 or L5 lie within the AC domain since when any of the loops are truncated, OmpT digests this domain releasing an ∼66 kDa α^43^ fragment into the extracellular milieu. We propose that the most C-terminal portion of the unfolded AC domain hydrogen bonds with the first β-strand in L4, forming a long β-sheet that also includes the L5 β-hairpin, thereby rapidly folding the AC domain and allowing processive in-register stacking of the β-helical rungs of the α^43^ domain to form the elongated β-helix (Fig. 7a). Our observation that the AC domain secreted by the β-barrel where all amino acids in the first β-strand in L5 were mutated to glycine had a more severe folding defect than the AC domain secreted by the β-barrel where all amino acids in the second β-strand in L5 were mutated to glycine suggests that in Ag43, the length of the templating β-sheet composed by L4 and L5 is important for efficient folding of the AC domain. In this scenario, the stretch of glycine residues in the first β-strand in L5 would break the four-stranded β-sheet composed by L4 and L5, leaving only the two-stranded L4 β-hairpin intact to template the folding of the AC domain β-hairpin (Fig. 7b). In contrast, the stretch of glycine residues in the second β-strand in L5 would leave a three-stranded β-hairpin to template the folding of the AC domain β-hairpin (Fig. 7c). We speculate that less of the ∼66 kDa OmpT-released α^43^ fragment is released from Ag43L5β2G than Ag43L5β1G because the longer length of the templating β-sheet provides greater stability to the L4-L5-AC β-hairpin interactions, thereby accelerating folding of the AC domain (albeit not as rapidly as in wild type Ag43) such that autocatalysis of α^43^ from β^43^ occurs faster than OmpT cleavage in the majority of Ag43L5β2G molecules. An interaction between the AC domain and the extracellular loops of the β-barrel domain as a prerequisite for the rapid folding of this region and the passenger domain accounts for the results of previous studies showing that mutations within the AC domain substantially perturb passenger folding^15–17^.

**Figure 7.**
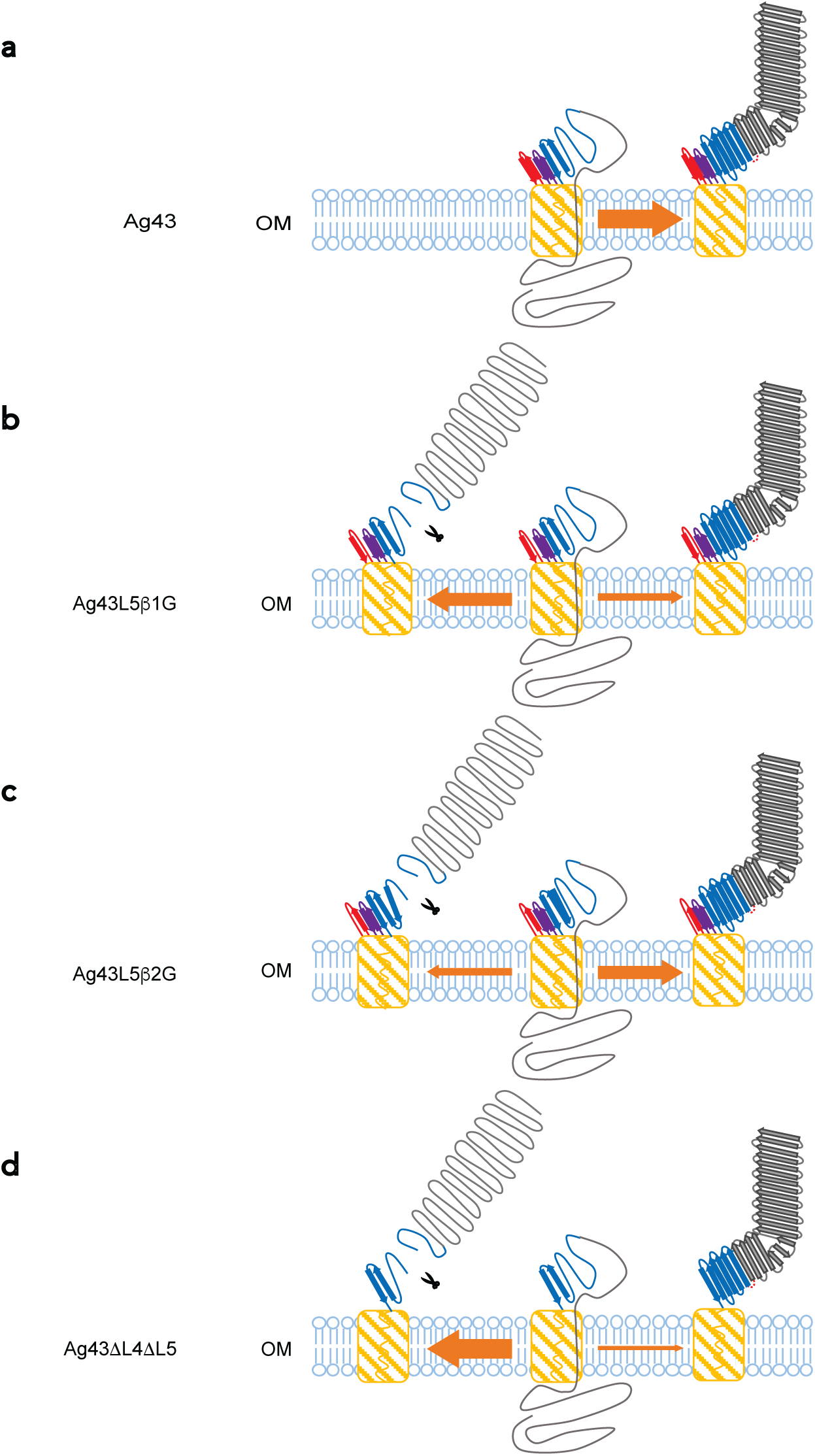
Proposed mechanism for Ag43a AC and passenger domain folding. **a** Schematic showing that the first β-strand in L4 (purple) interacts with the most C-terminal end of the AC domain (blue) during translocation, thereby rapidly nucleating processive folding and enabling in-register stacking of the passenger β-helical rungs (grey) to form the elongated fully folded β-helix. Once the extreme N-terminus of the α^43^ domain has traversed the β-barrel pore (yellow), autocatalysis liberates the α^43^ domain from the β^43^ domain (not shown), yet the two domains remain non-covalently associated. **b** Schematic shows that when all amino acids in the first β-strand in L5 (red) are mutated to glycine, the templating β-sheet is reduced to the L4 β-hairpin (purple), which slows folding of the AC domain such that OmpT cleavage occurs faster (thick orange arrow) than autocatalysis of α^43^ from β^43^ (thin orange arrow) in the majority of Ag43L5β1G molecules. **c** Schematic shows that when all amino acids in the second β-strand in L5 (red) are mutated to glycine, the templating β-sheet is reduced to the L4 β-hairpin (purple) and the first β-strand in L5 (red), which slows folding of the AC domain such that autocatalysis of α^43^ from β^43^ (thick orange arrow) occurs faster than OmpT cleavage (thin orange arrow) in the majority of Ag43L5β2G molecules. **d** Schematic shows that in the absence of the L4 and L5 β-hairpins, the potential for β-strand augmentation is completely abolished and interactions between the most C-terminal portion of the AC domain and the first β-strand in L4 is prevented, thereby causing a severe delay in folding of the AC and passenger domains. Because of the severe delay in AC and α^43^ domain folding, OmpT cleavage occurs much faster (thick orange arrow) than autocatalysis of α^43^ from β^43^ (thin orange arrow) in the majority of Ag431′L41′L5 molecules. In all cases, OmpT cleavage is indicated by scissors. BamA and other cellular machinery have been omitted for simplicity.

L4 and L5 appear to exhibit functional redundancy, allowing each β-hairpin to compensate in the absence of the other. This is supported by our observation that the α^43^ domain secreted by the double loop-truncated β-barrel has a substantially greater defect in folding *in vitro* and in mediating aggregation of *E. coli* MS4271′*ompT* compared to the α^43^ domains secreted by the single loop-truncated β-barrels. Our finding that the α^43^ domain released by the L4-truncated β-barrel shows a greater defect in folding *in vitro* than that released by the L5-truncated β-barrel suggests that L4 is the preferred templating β-hairpin. However, a greater dependence on L4 for folding of the α^43^ domain was not reflected *in vivo* where the α^43^ domains secreted by the L4- and- L5-truncated β-barrels showed a similar defect in aggregation of *E. coli* MS4271′*ompT*. When the potential for β-strand augmentation is completely abolished, we propose that hydrogen bonding between the most C-terminal portion of the unfolded AC domain and the first β-strand in L4 is prevented, and it is the absence of these initial interactions that severely delays folding of the AC domain and subsequent nucleation of α^43^ into a stack of in-register β-helical rungs. Consequently, OmpT can cleave the AC domain prior to autocatalysis of α^43^ from β^43^, resulting in the release of the ∼66 kDa α^43^ fragment into the extracellular milieu where it is unable to mediate its virulence function as a secreted protein (Fig. 7d).

We previously reported that truncation of L5 in the Pet β-barrel domain results in complete digestion of the passenger domain by exogenous proteases in bacterial cells and *in vitro*^34^. This contrasts with Ag43 where we observe delayed folding of the AC and α^43^ domains secreted by the loop-truncated β-barrels. In bacterial cells, delayed folding of the AC domains secreted by the loop-truncated β-barrels results in their digestion by endogenous OmpT. We propose that the slow-folding of the AC domains at the bacterial cell surface results in the exposure of OmpT-digestion sites that are not exposed in the AC domain secreted by the wild-type β-barrel because it is rapidly folded, thereby leaving the slow-folding AC domains susceptible to digestion by OmpT. We believe it unlikely that the distinct requirements of the long extracellular β-barrel loops for the folding of Pet versus Ag43 could be due to their distinct structures; while the long right-handed β-helix is common to both proteins, the Pet passenger domain contains a N-terminal chymotrypsin-like subdomain (conserved in all SPATEs) that projects from the β-helix to cleave fodrin, resulting in cytoskeletal disruption and host cell toxicity^48^. This is because folding of all passenger domains into β-helices in bacterial cells presumably occurs via the same vectorial pathway, whereby passenger folding is hypothesized to be coupled to translocation such that folding starts at the C-terminus (i.e., at the AC domain) and propagates through to the N-terminus as additional unfolded segments of the passenger domain emerge from the β-barrel. A plausible reason for this difference could be because L4 and L5 appear functionally redundant in Ag43, but not in Pet where only L5 is essential for passenger folding and L4 is required for efficient translocation of the passenger domain to the bacterial cell surface^34^. Certainly, the AC and α^43^ domains secreted by the double loop-truncated β-barrel were only slightly less susceptible to trypsinolysis than the Pet passenger domain secreted from the L5-truncated β-barrel^34^.

Therefore, the question becomes is there a functional value to having two extracellular loops that function to mediate passenger folding when one extracellular loop seems to suffice in the case of Pet? We wonder if the answer to this question has to do with the location of the AC domains post autocatalysis and the passengers’ biological functions. After processing of SPATEs via an autocatalytic intra-barrel cleavage reaction, the AC domain remains attached to the passenger domain^49–51^, in the case of Pet, resulting in the release of the passenger domain into the extracellular milieu so that it can enter epithelial cells to cause cellular toxicity^48^. The AC domain is not required for either of these processes. This contrasts with Ag43a where the AC domain remains attached to the membrane-embedded β-barrel domain, and α^43^ remains non-covalently associated with β^43^ to mediate bacterial aggregation despite its autocatalytic processing. Given that α^43^ remains associated with the single- and- double-loop-truncated translocation domains after refolding and autocatalysis *in vitro*, it is unlikely that L4 and L5 are involved in anchoring the α^43^ domain to the bacterial surface. Nevertheless, it is tempting to speculate a role for the long extracellular loops of the β-barrel domain post passenger translocation and folding. Given that L4, L5, and the AC domain are in close spatial proximity, it is conceivable that these loops remain in contact with the AC domain to stabilize the folded state of this region and the α^43^ β-helix, and/or to maintain the positioning of the elongated β-helix relative to the cell surface. In both scenarios, this would allow α^43^ to efficiently carry out its biological function. Here, the absence of one extracellular loop would be less detrimental than the absence of both extracellular loops. This fits well with the small delay in aggregation observed for *E. coli* MS427Δ*ompT* expressing Ag43ΔL4 or Ag43ΔL5 relative to that harboring wild type Ag43, and with the failure of bacteria expressing Ag43ΔL4ΔL5 to mediate any significant cell aggregation. Nevertheless, the increasing aggregative ability of *E. coli* MS427Δ*ompT* expressing Ag43ΔL4, Ag43ΔL5, or Ag43ΔL4ΔL5 over the time-course of the aggregation assay does suggest that the compromised biological function of these mutant proteins is also a consequence of delayed folding in a population of α^43^ molecules.

## Materials and Methods

### Reagents and bacterial strains

Unless otherwise stated, bacteria were grown at 37 °C in Luria–Bertani broth and where necessary, supplemented with 100 μg mL^−1^ ampicillin and/or 30 μg mL^−1^ kanamycin and, 0.2% (wt/vol) _L_-arabinose or 0.5 mM isopropylthio-β-galactoside (IPTG). Details of the *E. coli* strains [MS427, MS4271′*ompT*, and BL21 (DE3)] and plasmids used in this study are presented in Supplementary Table 1. Primers used for plasmid construction are presented in Supplementary Table 2.

### Construction of an *E. coli* MS427 *ompT* mutant

The 11 Red recombinase system^52^ was used to construct an in-frame *ompT* knock-out mutant of *E. coli* MS427.

### Plasmid construction

pCO4 (called pAg43 in this current study) has been described previously^38^. To construct pAg43ΔL4 and pAg43ΔL5, the coding sequence for the predicted loops L4 and L5, respectively of the Ag43a β-barrel domain were deleted from the full-length *flu* (i.e., *ag43a*) gene in pAg43 (locus tag c3655) from *E. coli* CFT073^36^, and two extra G residues were introduced to provide flexibility. pAg43ΔL4, pAg43ΔL5, and pAg43L5β1G were constructed from pAg43 by Epoch Life Science (USA) and GenScript, respectively. *Sex*AI-*Afe*I fragments comprising nucleotide sequence predicted to code for Ag43a with truncations in L4 and L5, and with glycine residues in place of predicted β2 in L5 were synthesized *de novo* by Integrated DNA Technologies (IDT), amplified using primers Ag43SexAIFw and Ag43AfeIRv, and cloned into pAg43, pre-digested with the same restriction enzymes, to create pAg43ΔL4ΔL5 and pAg43β2/G. To construct pAg43ΔL4ΔL5-AB, megaprimer PCR was performed as described previously^13,32,34^. Briefly, the round 1 PCR was performed using 500 ng of template DNA (pAg43) with 1 μg mL^−1^ each of primers MPAg43D^552^AFw and Ag43SexAIRv. The round 2 PCR was performed with 4 μg of megaprimer and 1 μg of primer Ag43KpnIFw using 500 ng of template DNA (pAg43). The resulting amplicon and target vectors (pAg43 and pAg43ΔL4ΔL5) were then digested with KpnI and SexAI and ligated. To construct pAg43^Δ1-206^, pAg43ΔL5^Δ1-206^, pAg43ΔL4^Δ1-206^, and pAg43ΔL4ΔL5^Δ1-206^, the Ag43a β-barrel domain, AC domain, and last 346 residues of the Ag43a passenger domain were amplified using 500 ng of template DNA (pAg43, pAg43ΔL4, pAg43ΔL5, and pAg43ΔL4L5) with 1 mg mL^−1^ each of primers NdeIAg43PD346Fw and XhoIAg43bRvHis. To construct pAg43^Δ1-705^, pAg43ΔL5^Δ1-705^, and pAg43ΔL4^Δ1-705^, and pAg43ΔL4L5^Δ1-705^ the Ag43a β-barrel domain was amplified using 500 ng of template DNA (pAg43, pAg43ΔL4, pAg43ΔL5, and pAg43ΔL4L5) with 1 μg/mL each of primers NdeIAg43bFw and XhoIAg43bRvHis. The subsequent amplicons and target vector (pET22b+) were then digested with *Nde*I and *Xho*I, and ligated for an in-frame C-terminal hexahistidine (his)-tag fusion. All DNA modifications were confirmed by sequencing.

### Immunodetection of the Ag43 β-domain

To obtain antibodies specific for the Ag43 β-barrel domain, Ag43^Δ1-705^ was refolded *in vitro* as described below and concentrated to ∼1 mg mL^−1^ using a centrifugal filter with a 30 kDa molecular weight cut-off. Rabbits were immunized 3 times with 200 μg of purified recombinant Ag43^Δ1-705^ at 4-week intervals at the Walter and Eliza Hall Institute of Medical Research (WEHI) Antibody Facility. Rabbits were bled on Day 68 and the resulting antiserum was protein-A purified by the WEHI Antibody Facility.

### *In vivo* protein expression assays

In all cases, bacterial cultures were grown to an OD_600_ between 0.5-0.6 prior to addition of 0.2 % arabinose for 2 hours to induce Ag43 expression. For proteinase K assays of whole cells, three 1 mL aliquots were removed from each culture. The first aliquot was added to a tube containing trichloroacetic acid (TCA) at a final concentration of 10% (wt/vol) and placed on ice. The second aliquot was added to a tube containing proteinase K at a final concentration of 200 μg mL^−1^ and incubated on ice for 20 minutes to digest Ag43 exposed on the cell surface. 1 mM phenylmethanesulfonyl fluoride (PMSF) was added to stop the protease reaction prior to TCA precipitation. All TCA precipitated samples were washed 2 times with acetone, dried, and resuspended in 100 mL of sodium dodecyl sulphate polyacrylamide gel electrophoresis (SDS-PAGE) loading buffer per OD_600_ units of cells (third aliquot). 5 μL of the normalized samples were separated by SDS-PAGE on 12 % gels and visualized by immunoblotting using anti-α^43^ and anti-Ag43a β-barrel antibodies at 1:200,000 and 1:10,000 dilutions, respectively. The anti-α^43^ antibody has been described and validated previously^36^.

For proteinase K assays of culture supernatant proteins, the OD_600_ of cultures were measured and pelleted (4,500 x *g*, 5 min, 4 °C) prior to filtering of the supernatant fractions through 0.22 μm pore-size filters. A 10% (wt/vol) final concentration of TCA was then used to precipitate Ag43 released into the culture supernatant as described above. For proteinase K time course assays of culture supernatant proteins, the OD_600_ of cultures were measured and supernatant fractions isolated as described above. Two 1 mL aliquots were removed from the supernatant fractions at increasing time points. The first aliquot was added to a tube containing a 10% (wt/vol) final concentration of TCA and placed on ice. The second aliquot was added to a tube containing a 200 μg mL^−1^ concentration of proteinase K and incubated on ice for 20 minutes to digest Ag43a released into the culture supernatant. The protease reaction was stopped by the addition of 1 mM PMSF prior to TCA precipitation. In all cases, samples were separated by 12% SDS-PAGE and subjected to immunoblotting using anti-α^43^ antibody at a 1:200,000 dilution as described above.

To demonstrate that the amount of PK in use did not permeabilize the outer membrane of *E. coli* MS427 or *E. coli* MS427Δ*ompT*, antibodies against the periplasmic chaperone SurA, was used for immunoblotting on the same samples to show that SurA was protected from PK digestion in all assays (Supplementary Fig. 5, left and right panels, respectively).

### Heat modifiability assay

After protein expression, cells were harvested and total membranes were isolated by ultracentrifugation as described previously^32,34^. Briefly, 10 μg of total membranes were mixed with SDS-PAGE loading buffer and then heated at 25 ℃ or 100 ℃ for 10 minutes. Samples were resolved by SDS-PAGE on 5-14 % gradient gels and subjected to immunoblotting using anti-Ag43 β-barrel domain antibodies as described above.

### Cell aggregation assay

Assays for cell-aggregation were performed as described previously with minor modifications^36^. Briefly, Ag43, Ag43ΔL4, Ag43ΔL5, or Ag43ΔL4ΔL5 were expressed as described above and adjusted to equivalent OD_600_ of 2.0, shaken vigorously, and then left to settle at room temperature. Three 100 μL samples were taken approximately 0.2 cm from the top of each culture, and the OD_600_ was measured at the zero time point and thereafter at 1 h intervals for 5 h.

### Protein refolding and purification

Protein expression and refolding were carried out as previously described with minor modifications^32,34^. *E. coli* BL21 (DE3) cells were transformed with pAg43^Δ1-206^, pAg43ΔL5^Δ1-206^, pAg43ΔL4^Δ1-206^, pAg43ΔL4ΔL5^Δ1-206^, pAg43^Δ1-705^, pAg43ΔL4^Δ1-705^, pAg43ΔL5^Δ1-705^, or pAg43ΔL4ΔL5^Δ1-705^, grown to OD_600_ = 0.3, and induced with 0.5 mM IPTG for 4 hours at 37 ℃. Cells were harvested (6,000 x *g*, 15 min, 4 ℃), resuspended in 50 mM tris pH 8.0, 150 mM NaCl, 1% Triton X-100 (wt/vol), incubated with 125 μg mL^−1^ lysozyme for 10 minutes on ice followed by 10 μg mL^−1^ DNase and 5 mM MgCl_2_ for 30 minutes on ice prior to lysis via EmulsiFlex (15,000 pounds per square inch). Inclusion bodies were collected by centrifugation (10,000 x *g*, 10 min, 4 ℃), washed three times with 50 mM tris pH 8.0, 150 mM NaCl, 1 % Triton X-100 (wt/vol) and once with 50 mM tris pH 8.0, 150 mM NaCl, and solubilized in 50 mM tris pH 8.0, 8 M urea. Inclusion bodies were normalized to 1 mg mL^−1^ and refolded by a rapid 10-fold dilution of the urea into 50 mM tris pH 8.0, 150 mM NaCl, 0.5 % (w/v) lauryldimethylamine oxide (LDAO) at 37 ℃. Affinity purification using HisTrap^TM^ HP columns was used to try and separate the α^43^ and β^43^ domains after folding and cleavage of the Ag43^Δ1-206^, Ag43ΔL5^Δ1-206^, Ag43ΔL4^Δ1-206^, and Ag43ΔL4ΔL5^Δ1-206^ proteins, which ended up co-eluting.

### N-terminal sequencing

For N-terminal amino acid sequencing of β^43Δ1-206^ and the ∼66 kDa OmpT-released α^43^ fragment of Ag43ΔL5, proteins were separated by 12 % SDS-PAGE and transferred onto a polyvinylidene difluoride (PVDF) sequencing-grade membrane. Proteins were visualized by staining with Coomassie brilliant blue (R-250), and the bands corresponding to protein fragments of interest were excised and sequenced by automated Edman degradation at the Monash University Biomedical Proteomics Facility.

### Limited trypsinolysis

Kinetics of trypsin cleavage profiles during Ag43^Δ1-206^, Ag43ΔL5^Δ1-206^, Ag43ΔL4^Δ1-206^, Ag43ΔL4ΔL5^Δ1-206^, Ag43^Δ1-705^, Ag43ΔL4^Δ1-705^, Ag43ΔL5^Δ1-705^, and Ag43ΔL4ΔL5^Δ1-705^ refolding were initiated as described above and followed by limited treatment with 20 μg mL^−1^ trypsin for 20 minutes on ice, as previously described^32,34^. The digestion and folding reactions were quenched with 1 mM PMSF and SDS loading buffer, respectively. The resulting trypsin-resistant fragments were resolved by 12% SDS-PAGE and visualized by immunoblotting with anti-α^43^ and anti-Ag43a β-barrel domain antibodies as described above.

### Biophysical analysis of protein structure

Far-UV CD spectra were collected from 200 to 260 nm using a Chirascan spectrometer as previously described^34^. Briefly, samples containing 0.1 mg mL^−1^ of refolded protein were measured with a 1 mm cuvette, 1.0 nm bandwidth, 1 s integration time, and 20 nm/min scanning speed at room temperature. The spectra were corrected for buffer contribution, and three scans were averaged per sample. For thermal denaturation measurements, the denaturation of β-sheet structure of samples containing 0.1 mg mL^−1^ of refolded protein was monitored at 218 nm as a function of temperature from 25 ℃ to 94 ℃ at a temperature change of 1 ℃ min^−1^ as described previously^34^. In all cases, the CD spectra were normalized to the mean residue molar ellipticity.

### Generation of Ag43 β-barrel model

AlphaFold3 (https://alphafoldserver.com/) was used to predict the structure of the Ag43a translocator domain (P^553^-F^1040^). The structural model has a predicted template modelling (pTM) value of 0.79 (pTM >0.5 indicates high confidence). PyMOL (https://pymol.org/2/) was used to generate structural figures.

## Data availability

All data generated and analysed during this study are included in this published article and its Supplementary Information files. Other data are available from the corresponding author upon reasonable request.

## Acknowledgements

We thank Fiona Lewis for technical assistance, John Carver for access to the Chirascan spectrometer, as well as Kevin Saliba for critical comments on the manuscript. This work was supported by the Australian Research Council (ARC) Discovery Project grant DP160103294 (to D.L.L. and I.R.H.), Future Fellowship FT150100452 (to D.L.L), and a National Health and Medical Research Council (NHMRC) Project grant GNT1143638 (to D.L.L. and B.H).

## Author contributions

X.Y. and D.L.L. conceived and designed the experiments. X.Y., M.D.J., J.L., J.Z., A.W.L., J.J.P., and M-D.P., performed the experiments. X.Y., B.H., M.A.S., G.H.M.H., I.R.H., M.T.D. and D.L.L. analyzed the data. X.Y., M.T.D. and D.L.L. wrote the manuscript. All authors have reviewed the manuscript.

## References

1 Celik, N. et al. A bioinformatic strategy for the detection, classification and analysis of bacterial autotransporters. PLoS One 7, e43245 (2012). 10.1371/journal.pone.0043245

2 Clarke, K. R. et al. Phylogenetic classification and functional review of autotransporters. Front. Immunol. 13, 921272 (2022). 10.3389/fimmu.2022.921272

3 Meuskens, I., Saragliadis, A., Leo, J. C. & Linke, D. Type V Secretion Systems: an overview of passenger domain functions. Front. Microbiol. 10, 1163 (2019). 10.3389/fmicb.2019.01163

4 Fan, E., Chauhan, N., Udatha, D. B. R. K. G., Leo, J. C. & Linke, D. Type V secretion systems in bacteria. Microbiol. Spectr. 4, 0.1128/microbiolspec.VMBF-0009-2015 (2016). doi:10.1128/microbiolspec.VMBF-0009-2015

5 Pohlner, J., Halter, R., Beyreuther, K. & Meyer, T. F. Gene structure and extracellular secretion of *Neisseria gonorrhoeae* IgA protease. Nature 325, 458–462 (1987). 10.1038/325458a0

6 Saurí, A. et al. Autotransporter β-domains have a specific function in protein secretion beyond outer-membrane targeting. J. Mol. Biol. 412, 553–567 (2011). 10.1016/j.jmb.2011.07.035

7 Doyle, M. T. & Bernstein, H. D. BamA forms a translocation channel for polypeptide export across the bacterial outer membrane. Mol. Cell. 81, 2000–2012.e2003 (2021). 10.1016/j.molcel.2021.02.023

8 Doyle, M. T. et al. Cryo-EM structures reveal multiple stages of bacterial outer membrane protein folding. Cell 185, 1143–1156.e1113. (2022). 10.1016/j.cell.2022.02.016

9 Junker, M. et al. Pertactin β-helix folding mechanism suggests common themes for the secretion and folding of autotransporter proteins. Proc. Natl. Acad. Sci. U S A. 103, 4918–4923 (2006). 10.1073/pnas.0507923103

10 Peterson, J. H., Tian, P., Ieva, R., Dautin, N. & Bernstein, H. D. Secretion of a bacterial virulence factor is driven by the folding of a C-terminal segment. Proc. Natl. Acad. Sci. U S A. 107, 17739–17744 (2010). 10.1073/pnas.1009491107

11 Junker, M., Besingi, R. N. & Clark, P. L. Vectorial transport and folding of an autotransporter virulence protein during outer membrane secretion. Mol. Microbiol. 71, 1323–1332 (2009). 10.1111/j.1365-2958.2009.06607.x

12 Renn, J. P. & Clark, P. L. A conserved stable core structure in the passenger domain β-helix of autotransporter virulence proteins. Biopolymers 89, 420–427 (2008). 10.1002/bip.20924

13 Leyton, D. L. et al. Size and conformation limits to secretion of disulfide-bonded loops in autotransporter proteins. J. Biol. Chem. 286, 42283–42291 (2011). 10.1074/jbc.M111.306118

14 Vo, J. L. et al. Variation of Antigen 43 self-association modulates bacterial compacting within aggregates and biofilms. NPJ Biofilms Microbiomes 8, 20 (2022). 10.1038/s41522-022-00284-1

15 Ohnishi, Y., Nishiyama, M., Horinouchi, S. & Beppu, T. Involvement of the COOH-terminal pro-sequence of *Serratia marcescens* serine protease in the folding of the mature enzyme. J. Biol. Chem. 269, 32800–32806 (1994). 10.1016/S0021-9258(20)30062-4

16 Oliver, D. C., Huang, G., Nodel, E., Pleasance, S. & Fernandez, R. C. A conserved region within the *Bordetella pertussis* autotransporter BrkA is necessary for folding of its passenger domain. Mol. Microbiol. 47, 1367–1383 (2003). 10.1046/j.1365-2958.2003.03377.x

17 Dutta, P. R., Sui, B. Q. & Nataro, J. P. Structure-function analysis of the Enteroaggregative *Escherichia coli* plasmid-encoded toxin autotransporter using scanning linker mutagenesis. J. Biol. Chem. 278, 39912–39920 (2003). 10.1074/jbc.M303595200

18 Kajava, A. V. & Steven, A. C. The turn of the screw: Variations of the abundant β-solenoid motif in passenger domains of Type V secretory proteins. J. Struct. Biol. 155, 306–315 (2006). 10.1016/j.jsb.2006.01.015

19 Leyton, D. L., Rossiter, A. E. & Henderson, I. R. From self sufficiency to dependence: mechanisms and factors important for autotransporter biogenesis. Nat. Rev. Microbiol. 10, 213–225 (2012). 10.1038/nrmicro2733

20 Ieva, R., Tian, P., Peterson, J. H. & Bernstein, H. D. Sequential and spatially restricted interactions of assembly factors with an autotransporter β domain. Proc. Natl. Acad. Sci. U S A. 108, E383–E391 (2011). 10.1073/pnas.1103827108

21 Ieva, R. & Bernstein, H. D. Interaction of an autotransporter passenger domain with BamA during its translocation across the bacterial outer membrane. Proc. Natl. Acad. Sci. U S A. 106, 19120–19125 (2009). 10.1073/pnas.0907912106

22 Selkrig, J. et al. Discovery of an archetypal protein transport system in bacterial outer membranes. Nat. Struct. Mol. Biol. 19, 506–510 (2012). 10.1038/nsmb.2261

23 Shen, H.-H. et al. Reconstitution of a nanomachine driving the assembly of proteins into bacterial outer membranes. Nat. Commun. 5, 5078 (2014). 10.1038/ncomms6078

24 Gruss, F. et al. The structural basis of autotransporter translocation by TamA. Nat. Struct. Mol. Biol. 20, 1318–1320 (2013). 10.1038/nsmb.2689

25 Ruiz-Perez, F. et al. Roles of periplasmic chaperone proteins in the biogenesis of serine protease autotransporters of *Enterobacteriaceae*. J. Bacteriol. 191, 6571–6583 (2009). 10.1128/jb.00754-09

26 Pavlova, O., Peterson, J. H., Ieva, R. & Bernstein, H. D. Mechanistic link between β barrel assembly and the initiation of autotransporter secretion. Proc. Natl. Acad. Sci. U S A. 110, E938–E947 (2013). 10.1073/pnas.1219076110

27 Doyle, M. T. & Bernstein, H. D. Bacterial outer membrane proteins assemble via asymmetric interactions with the BamA β-barrel. Nat. Commun. 10, 3358–3358 (2019). 10.1038/s41467-019-11230-9

28 Voulhoux, R., Bos, M. P., Geurtsen, J., Mols, M. & Tommassen, J. Role of a highly conserved bacterial protein in outer membrane protein assembly. Science 299, 262–265 (2003). 10.1126/science.1078973

29 Malinverni, J. C. et al. YfiO stabilizes the YaeT complex and is essential for outer membrane protein assembly in *Escherichia coli*. Mol. Microbiol. 61, 151–164 (2006). 10.1111/j.1365-2958.2006.05211.x

30 Wu, T. et al. Identification of a multicomponent complex required for outer membrane biogenesis in *Escherichia coli*. Cell 121, 235–245 (2005). 10.1016/j.cell.2005.02.015

31 Shen, C. et al. Structural basis of BAM-mediated outer membrane β-barrel protein assembly. Nature 617, 185–193 (2023). 10.1038/s41586-023-05988-8

32 Leyton, D. L. et al. A mortise–tenon joint in the transmembrane domain modulates autotransporter assembly into bacterial outer membranes. Nat. Commun. 5, 4239 (2014). 10.1038/ncomms5239

33 Zhai, Y. et al. Autotransporter passenger domain secretion requires a hydrophobic cavity at the extracellular entrance of the β-domain pore. Biochem. J. 435, 577–587 (2011). 10.1042/bj20101548

34 Yuan, X. et al. Molecular basis for the folding of β-helical autotransporter passenger domains. Nat. Commun. 9, 1395 (2018). 10.1038/s41467-018-03593-2

35 Ageorges, V. et al. Differential homotypic and heterotypic interactions of antigen 43 (Ag43) variants in autotransporter-mediated bacterial autoaggregation. Sci. Rep. 9, 11100 (2019). 10.1038/s41598-019-47608-4

36 Heras, B. et al. The antigen 43 structure reveals a molecular Velcro-like mechanism of autotransporter-mediated bacterial clumping. Proc. Natl. Acad. Sci. U S A. 111, 457–462 (2014). 10.1073/pnas.1311592111

37 Klemm, P., Hjerrild, L., Gjermansen, M. & Schembri, M. A. Structure-function analysis of the self-recognizing Antigen 43 autotransporter protein from *Escherichia coli*. Mol. Microbiol. 51, 283–296 (2004). 10.1046/j.1365-2958.2003.03833.x

38 Ulett, G. C. et al. Functional analysis of Antigen 43 in uropathogenic *Escherichia coli* reveals a role in long-term persistence in the urinary tract. Infect. Immun. 75, 3233–3244 (2007). 10.1128/iai.01952-06

39 Rojas-Lopez, M. et al. Identification of the autochaperone domain in the Type Va Secretion System (T5aSS): prevalent feature of autotransporters with a β-helical passenger. Front. Microbiol. 8, 2607 (2018). 10.3389/fmicb.2017.02607

40 Abramson, J. et al. Accurate structure prediction of biomolecular interactions with AlphaFold 3. Nature 630, 493–500 (2024). 10.1038/s41586-024-07487-w

41 Wang, X., Nyenhuis, S. B. & Bernstein, H. D. The translocation assembly module (TAM) catalyzes the assembly of bacterial outer membrane proteins in vitro. Nat. Commun. 15, 7246 (2024). 10.1038/s41467-024-51628-8

42 Nakamura, K. & Mizushima, S. Effects of heating in dodecyl sulfate solution on the conformation and electrophoretic mobility of isolated major outer membrane proteins from *Escherichia coli* K-12. J. Biochem. 80, 1411–1422 (1976). 10.1093/oxfordjournals.jbchem.a131414

43 Xue, Z., Pang, Y. & Quan, S. Revisiting the functions of periplasmic chaperones in the quality control of the autotransporter Ag43 using a phenotypically homogeneous *Escherichia coli* strain. Biochem Biophys Res Commun 591, 37–43 (2022). 10.1016/j.bbrc.2021.12.110

44 Marie-Eve Charbonneau, J. J. & Mourez, M. Autoprocessing of the *Escherichia coli* AIDA-I autotransporter. J. Biol. Chem. 284, 17340–17351 (2009). 10.1074/jbc.M109.010108

45 Caffrey, P. & Owen, P. Purification and N-terminal sequence of the alpha subunit of antigen 43, a unique protein complex associated with the outer membrane of *Escherichia coli*. J. Bacteriol. 171, 3634–3640 (1989). 10.1128/jb.171.7.3634-3640.1989

46 Kjærgaard, K., Schembri, M. A., Ramos, C., Molin, S. & Klemm, P. Antigen 43 facilitates formation of multispecies biofilms. Environ. Microbiol. 2, 695–702 (2000). 10.1046/j.1462-2920.2000.00152.x

47 Imai, K. & Mitaku, S. Mechanisms of secondary structure breakers in soluble proteins. Biophysics (Nagoya-shi) 1, 55–65 (2005). 10.2142/biophysics.1.55

48 Canizalez-Roman, A. & Navarro-García, F. Fodrin CaM-binding domain cleavage by Pet from enteroaggregative *Escherichia coli* leads to actin cytoskeletal disruption. Mol. Microbiol. 48, 947–958 (2003). 10.1046/j.1365-2958.2003.03492.x

49 Barnard, T. J., Dautin, N., Lukacik, P., Bernstein, H. D. & Buchanan, S. K. Autotransporter structure reveals intra-barrel cleavage followed by conformational changes. Nat. Struct. Mol. Biol. 14, 1214–1220 (2007). http://www.nature.com/nsmb/journal/v14/n12/suppinfo/nsmb1322_S1.html

50 Barnard, T. J. et al. Molecular basis for the activation of a catalytic asparagine residue in a self-cleaving bacterial autotransporter. J. Mol. Biol. 415, 128–142 (2012). 10.1016/j.jmb.2011.10.049

51 Dautin, N., Barnard, T. J., Anderson, D. E. & Bernstein, H. D. Cleavage of a bacterial autotransporter by an evolutionarily convergent autocatalytic mechanism. EMBO J. 26, 1942–1952 (2007). http://www.nature.com/emboj/journal/v26/n7/suppinfo/7601638a_S1.html

52 Datsenko, K. A. & Wanner, B. L. One-step inactivation of chromosomal genes in *Escherichia coli* K-12 using PCR products. Proc. Natl. Acad. Sci. U S A. 97, 6640–6645 (2000). 10.1073/pnas.120163297

