## Supplementary Information for "Extracellular loops of the β-barrel domain catalyze rapid folding for function of self-associating autotransporters"

**a**

| Amino Acid Sequence | | | | | | | | | | |
|---|---|---|---|---|---|---|---|---|---|---|
| **Residue** | **1** | **2** | **3** | **4** | **5** | **6** | **7** | **8** | **9** | **10** |
| Major Sequence: | P | T | N | V | T | | | | | |
| Minor Sequence: | | | | | | | | | | |
| **Residue** | **11** | **12** | **13** | **14** | **15** | **16** | **17** | **18** | **19** | **20** |
| Major Sequence: | | | | | | | | | | |
| Minor Sequence: | | | | | | | | | | |

**b**

| Amino Acid Sequence | | | | | | | | | | |
|---|---|---|---|---|---|---|---|---|---|---|
| **Residue** | **1** | **2** | **3** | **4** | **5** | **6** | **7** | **8** | **9** | **10** |
| Major Sequence: | A | D | I | V | V | | | | | |
| Minor Sequence: | | V G | | | | | | | | |
| **Residue** | **11** | **12** | **13** | **14** | **15** | **16** | **17** | **18** | **19** | **20** |
| Major Sequence: | | | | | | | | | | |
| Minor Sequence: | | | | | | | | | | |

**Supplementary Figure 1. N-terminal amino acid sequencing using Edman degradation.** The amino acids at the beginning of the fragments corresponding to the refolded wild-type β-barrel ($\beta^{43\Delta 1\text{-}206}$) (**a**) and the ~66 kDa OmpT-released $\alpha^{43}$ species of Ag43ΔL5 (**b**) are numbered from 1 to 5.

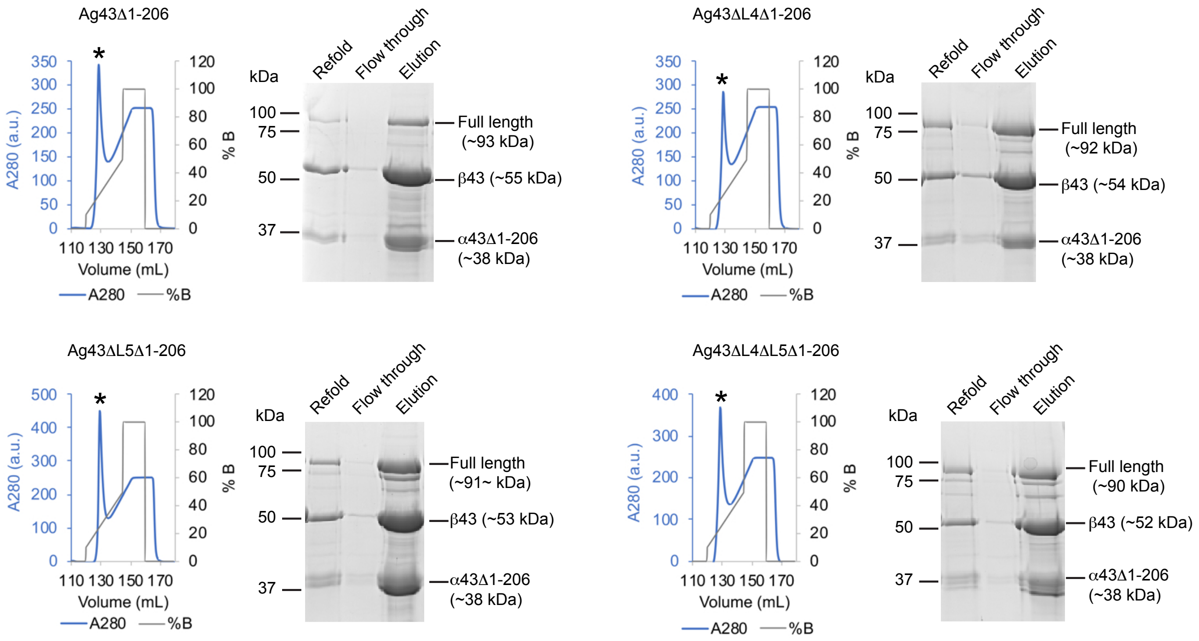


**Supplementary Figure 2. Co-elution of the $\alpha^{43}$ and $\beta^{43}$ domains.** Cleaved $Ag43^{\Delta 1\text{-}206}$, $Ag43\Delta L4^{\Delta 1\text{-}206}$, $Ag43\Delta L5^{\Delta 1\text{-}206}$, and $Ag43\Delta L4\Delta L5^{\Delta 1\text{-}206}$ were purified through immobilized metal affinity chromatography (IMAC) using Ni-NTA columns (* indicates their elution peak) (left panels), resulting in co-elution of the ~38 kDa $\alpha^{43}$ and ~55 kDa $\beta^{43}$ domains as analyzed by SDS-PAGE and Coomassie staining (right panels).

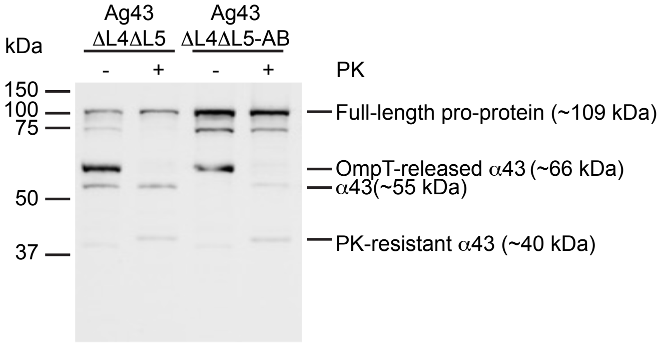


**Supplementary Figure 3. The ~55 kDa and ~40 kDa protease-resistant fragments arise from the ~66 kDa passenger fragment.** Expression of Ag43ΔL4ΔL5 and Ag43ΔL4ΔL5-AB (autocatalysis blocked; Ag43L4ΔL5D$^{552}$A), and sensitivity to proteinase K (PK) in *E. coli* MS427 monitored by Western immunoblotting with anti-α$^{43}$ antibody.

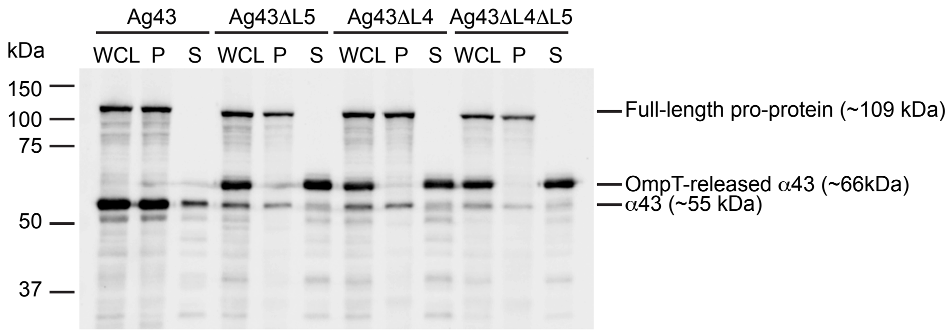


**Supplementary Figure 4. Localization of the ~66 kDa OmpT-released $\alpha^{43}$ fragment in the culture supernatant.** Fractionation of *E. coli* MS427 expressing Ag43, Ag43ΔL4, Ag43ΔL5, or Ag43ΔL4ΔL5 showing whole cell lysates (WCL), bacterial cell pellets (P), and culture supernatant (S) fractions. Samples were visualized by Western immunoblotting with anti-$\alpha^{43}$ antibody.

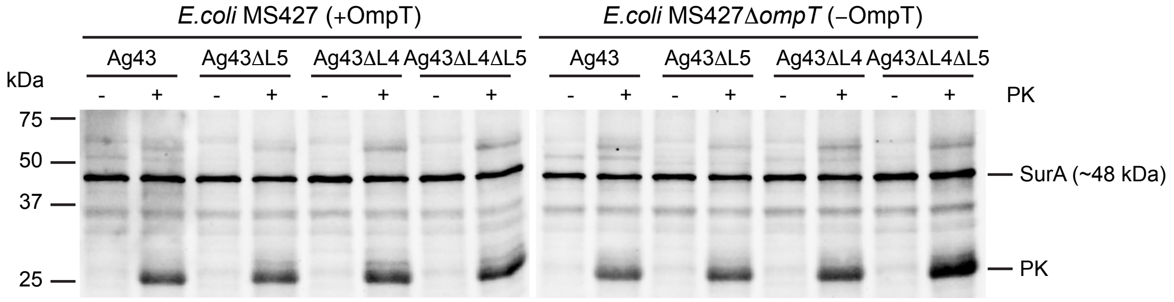


**Supplementary Figure 5. Proteinase K does not permeabilize the outer membrane of *E. coli* MS427 or *E. coli* MS427Δ*ompT*.** Ag43 expression in *E. coli* MS427 and *E. coli* MS427Δ*ompT*. All samples were analyzed by Western immunoblotting using antibody specific for the periplasmic protein, SurA. In both bacterial strains, the intact SurA protein was observed in −PK and +PK samples. Note that bands corresponding to the migration position of proteinase K are also visible. These data show that the concentration of proteinase K used does not permeabilize the *E. coli* outer membrane, which remains intact during proteinase K treatment.

**Supplementary Table 1. Strains and plasmids used in this study**

| Strain/Plasmid | Relevant description | Reference |
|---|---|---|
| *E. coli* MS427 | *ilvG*, *rfb-50*, *thi*, Δ*flu* (MG1655 Δ*flu*) | [1] |
| *E. coli* MS427 *ompT*::*kan*(Δ*ompT*) | An in-frame *ompT* knock-out mutant of *E. coli* MS427 | This study |
| *E. coli* BL21 (DE3) | F– *omp*T *hsd*SB(rB–, mB–) *gal dcm* (DE3) | Invitrogen |
| pBADMycHisA | Arabinose-inducible expression vector, ampicillin resistant | Invitrogen |
| pAg43 | pBADMycHisA derivative called pCO4 expressing Ag43 [*fluA* gene (c3655) from CFT073], ampicillin and kanamycin resistant | [2] |
| pAg43ΔL4 | pAg43 derivative expressing Ag43 with a L4 truncation | Epoch Life Sciences/ This study |
| pAg43ΔL5 | pAg43 derivative expressing Ag43 with a L5 truncation | Epoch Life Sciences/ This study |
| pAg43ΔL4ΔL5 | pAg43 derivative expressing Ag43 with a L4 and L5 double truncation | IDT/This study |
| pAg43L5β1G | pAg43 derivative expressing Ag43 where $D^{973}$, $M^{974}$, $R^{975}$, $V^{976}$ in β-strand 1 of L5 are mutated to G to assess if these residues play a role in passenger domain folding | GenScript/ This study |
| pAg43L5β2G | pAg43 derivative expressing Ag43 where $T^{986}$, $F^{987}$, $S^{988}$, $P^{989}$, in β-strand 2 of L5 are mutated to G to assess if these residues play a role in passenger domain folding | GenScript / This study |
| pAg43ΔL4ΔL5-AB | pAg43ΔL4ΔL5 derivative expressing Ag43ΔL4ΔL5 where $D^{552}$ in passenger is mutated to A to inhibit autocatalytic cleavage (AB, autocatalysis-blocked) | This study |
| pET-22b+ | IPTG-inducible expression vector allowing a C-terminal hexahistidine-tag fusion, ampicillin resistant | Novagen |
| pAg43$^{\Delta 1\text{-}206}$ | Truncated pAg43 derivative containing a hexahistidine-tagged β-barrel domain and the last 346 residues of the passenger domain located immediately N-terminal to the AC domain | This study |

| | | |
|---|---|---|
| pAg43ΔL4$^{\Delta 1\text{-}206}$ | Truncated pAg43ΔL4 derivative containing a hexahistidine-tagged β-barrel domain and the last 346 residues of the passenger domain located immediately N-terminal to the AC domain | This study |
| pAg43ΔL5$^{\Delta 1\text{-}206}$ | Truncated pAg43ΔL5 derivative containing a hexahistidine-tagged β-barrel domain and the last 346 residues of the passenger domain located immediately N-terminal to the AC domain | This study |
| pAg43ΔL4ΔL5$^{\Delta 1\text{-}206}$ | Truncated pAg43ΔL4ΔL5 derivative containing a hexahistidine-tagged β-barrel domain and the last 346 residues of the passenger domain located immediately N-terminal to the AC domain | This study |
| pAg43$^{\Delta 1\text{-}705}$ | Truncated pAg43 derivative containing a hexahistidine-tagged β-barrel domain | This study |
| pAg43ΔL4$^{\Delta 1\text{-}705}$ | Truncated pAg43ΔL4 derivative containing a hexahistidine-tagged β-barrel domain | This study |
| pAg43ΔL5$^{\Delta 1\text{-}705}$ | Truncated pAg43ΔL5 derivative containing a hexahistidine-tagged β-barrel domain | This study |
| pAg43ΔL4ΔL5$^{\Delta 1\text{-}705}$ | Truncated pAg43ΔL4ΔL5 derivative containing a hexahistidine-tagged β-barrel domain | This study |
| pKD46 | Donor of λRed genes to enable λRed-dependent recombination | [3] |

Note that the numbers next to the amino acid residues correspond to their position relative to the full-length Ag43 protein (from $M^1$ to $F^{1040}$).

**Supplementary Table 2. Primers used in this study**

| Primer | Sequence | Reference |
|---|---|---|
| NdeIAg43PD346 Fw | 5’-GGGAATTC**CATATG**GTCGGAGGGACGGC-3’ | This study |
| NdeIAg43bFw | 5’-GGGAATTC**CATATG**GCTTATCGTGCAGAAGTCC-3’ | This study |
| XhoIAg43bRvHis | 5’-CCG**CTCGAG***AGAGCC*GAAGGTCACATTCAGTGTG-3’ | This study |
| Ag43AfeIRv | 5’-CCTTC**AGCGCT**GCTGCC-3’ | This study |
| Ag43SexAIFw | 5’-CCTGA**ACCTGGT**GAACGC-3’ | This study |
| Ag43KpnIFw | 5’-GT**GGTACC**CGGAGCGAC-3’ | This study |
| Ag43SexAIRv | 5’-GCGTTC**ACCAGGT**TCAGG-3’ | This study |
| MPAg43D$^{552}$AFw | 5’GCACTGTGCTGAACGGTGCCATTGCGCCCACGAATG TCACTCTCGCCTC-3’ | This study |

Restriction enzyme sequences are in bold font, insertion sequences (Gly-Ser linker) are italicized, and mutated sequences are underlined.

### Supplementary References


1 Reisner, A., Haagensen, J. A., Schembri, M. A., Zechner, E. L. & Molin, S. Development and maturation of *Escherichia coli* K-12 biofilms. *Mol. Microbiol.* **48**, 933-946 (2003). https://doi.org/10.1046/j.1365-2958.2003.03490.x

2 Ulett, G. C. *et al.* Functional analysis of Antigen 43 in uropathogenic *Escherichia coli* reveals a role in long-term persistence in the urinary tract. *Infect. Immun.* **75**, 3233-3244 (2007). https://doi.org/10.1128/iai.01952-06

3 Datsenko, K. A. & Wanner, B. L. One-step inactivation of chromosomal genes in *Escherichia coli* K-12 using PCR products. *Proc. Natl. Acad. Sci. U S A.* **97**, 6640-6645 (2000). https://doi.org/10.1073/pnas.120163297
